# Modelling the collective mechanical regulation of the structure and morphology of epithelial cell layers

**DOI:** 10.1101/2021.08.30.458304

**Authors:** Hamid Khataee, Madeleine Fraser, Zoltan Neufeld

**Affiliations:** School of Mathematics and Physics, The University of Queensland, St. Lucia, Brisbane, QLD 4072, Australia

## Abstract

The morphology and function of epithelial sheets play an important role in healthy tissue development and cancer progression. The maintenance of structure of closely packed epithelial layers requires the coordination of various mechanical forces within the cells and others resulting from interactions with other cells and other tissues or substrates. However, a general model for the combination of mechanical properties which determine the cell shape and the overall structure of epithelial layers remains elusive. Here, we propose a computational model, based on the Cellular Potts Model, to study the interplay between mechanical properties of cells and dynamical transitions in epithelial structures and cell shapes. We map out phase diagrams as functions of cellular properties and the orientation of cell division. Monolayers of squamous, cuboidal, and columnar cells are found when the axis of cell proliferation is perpendicular to the substrate. Monolayer-to-multilayer transition is promoted via cell extrusion, depending on the mechanical properties of cells and the orientation of cell division. The results and model predictions are discussed in the context of experimental observations.

## 1 Introduction

Understanding the mechanisms of the development of various tissue morphologies is a major challenge in biology [1]. Epithelial cell layers are the simplest living tissues that line organs throughout the body [2] that play an important role in regulating embryo development and account for about 90% of all cancers [3]. Morphogenesis of organ systems is driven by the ability of cells to survive and proliferate [4, 5], primarily regulated by cell growth factors and cell-substrate adhesion [4, 6, 7].

For many adherent cells, cell proliferation can only occur on a substrate [8]. The substrate maintains a dynamic force balance between the cell and its microenvironment, and thus, the loss of substrate or its abnormal stiffness can results in aberrant cellular behaviours, e.g., breast tumor progression [9]. As feedback loops, cells sense the stiffness of their environment by pulling against the extracellular matrix, through integrin-extracellular matrix linkages, and/or neighbouring cells [4, 9]. This process is dependent on cell–substrate and cell-cell adhesion, as well as the contractility of cell cortex [9]. Therefore, both integrins and growth factor receptors use cytoplasmic signaling pathways to regulate cell cycle progression and growth [6]. It has been shown that the probability of cell proliferation increases with increasing substrate stiffness [10] and cell area [5]. Yet, it remains inconclusive how different forces and regulatory mechanisms within cells can affect proliferation orientation; reviewed in [11, 12].

Earlier theoretical studies on epithelial morphology have explored two-dimensional (2D) mechanical model of a tubular epithelium [13, 14], geometric patterning of apical junctions [15–18], shapes of cells and the buckling of cell monolayers [1, 19]. Although these models are based on the mechanical properties of cells, they were mostly restricted to monolayers. To model the dynamic processes involved in the formation of epithelial cell layers, models of epidermal homeostasis were proposed based on probabilistic rules associated to different types of cells [20–22]. However, these models do not consider the shape of the cells and the role of cellular mechanics in modelling the transition between monolayers to multilayers. Therefore, it remains elusive how the mechanical properties of cells and their interactions determine cell aspect ratios and the formation of mono- and multilayered epithelial structures. Further, the role of the orientation of the plane of cell division, in combination with mechanical properties of cells, in modelling collective tissue morphology has not been explored.

Here, we propose a simple computational model for analysing the development of collective epithelial morphology using the Cellular Potts Model (CPM) [23, 24]. CPM is a computational modelling framework that can represent the essential features of the real-world epithelial cell dynamics, and allows general predictions of the behaviour and morphology of cells [25, 26]. We simulate the transition of cell shapes and the formation of mono- and multilayered structures by altering various mechanical properties of identical proliferative cells.

## 2 Theoretical Model

To simulate collective morphology of cells emerging through their mechanical properties and interactions on a substrate, we use a two-dimensional CPM [23, 24] which represents a cross-section of cells on a substrate on a plane perpendicular to the substrate. The CPM is an on-lattice model which is computationally simpler than most off-lattice models, e.g., vertex model [27, 28] and has been used to capture essential realistic features of epithelial cell dynamics [25], e.g., the dynamics of cell migration on short microlanes [29], circular micropatterns [30], and in a confluent sheet expanding into a free region [26].

The cells are represented on a lattice, where each cell covers a set of connected lattice sites (or pixels) and each pixel can only be occupied by one cell at a time. Here, the lattice is a rectangular surface (480 *×* 195 pixels in the *x*- and *y*-dimensions, respectively), representing a cross-sectional view to epithelial cells placed on a substrate. The expansion and retraction of the cell boundaries are determined by minimising a phenomenological energy *E*, defined in terms of the area *A*_*σ*_ and perimeter *L*_*σ*_ of each cell *σ* of *N* cells (indices *σ* = 1, …, *N*) [17, 26, 31–33] as:

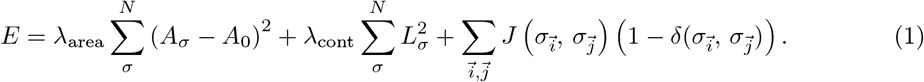

The first term models the compressibility of cells by penalising the deviation of cell areas from a target area *A*_0_. The second term represents the contractility of the cell cortex as a spring with zero equilibrium length (i.e., the target length of the cell perimeter is zero). The penalty parameter *λ*_cont_ represents cortical actomyosin contractility, around the lateral cell membrane [34]. The last term describes the cell-cell adhesion mediated by adhesion molecules, such as E-cadherin [35]. *J* is the boundary energy cost at neighbouring lattice sites 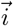 and 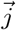. The Kronecker *δ* function prevents counting pixels that belong to the same cell. When both lattice sites 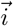 and 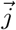 correspond to cells, 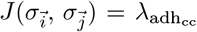; when one lattice site corresponds to cell and another site corresponds to the substrate 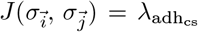; otherwise when one or both lattice sites represent empty space or boundary wall, the boundary energy cost *J* is set to zero. Note that 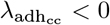 and 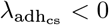 to represent that cells preferentially expand their boundaries shared with neighbouring cells or substrate. This is however balanced by the contractile tension along the cell cortex. The prefactors *λ*_area_, *λ*_cont_, and *λ*_adh_ reflect the relative importance of the corresponding cellular properties.

The dynamics of the CPM is defined by a stochastic series of elementary steps, where a cell expands or shrinks accommodated by a corresponding area change in the adjacent cell (or empty area) [24, 36]. The algorithm randomly selects two adjacent lattice sites 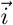 and 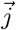, occupied by different cells 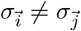. The elementary step is an attempt to copy 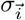 into the adjacent lattice site 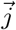, which takes place with probability

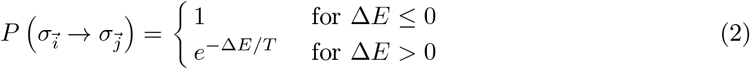

where Δ*E* is the change in functional (1) due to the elementary step considered, and the temperature parameter *T* is an arbitrary scaling factor. A Monte Carlo step (MCS) of the simulation, the natural unit of time in the model, is set to *n* elementary steps – where *n* is the total number of lattice sites in the simulated area [36]. Together, Equations (1, 2) imply that cell configurations which increase the penalties in functional (1) are less likely to occur. Thus, the cell population evolves through stochastic rearrangements in accordance with the biological dynamics incorporated into the effective energy function *E*.

Among multiple environmental factors that can regulate cell proliferation, cell growth factors and cell-substrate adhesion are most crucial [6, 7]: the probability of cell proliferation for individual cells increases with the cell area [5] and substrate stiffness [10]. We therefore define cell proliferation probability as a function of cell area and adhesion to the substrate in the form of the Hill function, which is widely used in mathematical modelling of binding of molecular structures of cells [37]. At every MCS, if a cell *σ* reaches its target area (i.e., *A*_*σ*_ *≥ A*_0_), the probability of proliferation is given by the following expression:

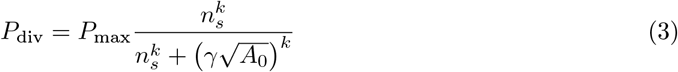

where *P*_max_ is the maximum probability of proliferation and *n*_*s*_ denotes the number of boundary pixels of a cell adjacent to the substrate, representing cell-substrate adhesion sites [38]. We assume that the Hill half-saturation threshold is given by the dimension in pixels of a square shaped cell, i.e., 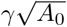 with a multiplicative factor *γ*. Together, Equation (3) expresses that the proliferation probability of a cell increases as the cell is more adhesive to the substrate. This is consistent with experiments [4, 9, 10] where increased area of cell-substrate contact enhanced cell growth, and thus proliferation. Further, *P*_div_ = 0 for cells not adhered to the substrate, representing that cell proliferation can occur only on the substrate [8].

Our simulations are implemented using the open-source software package CompuCell3D (CC3D) [36]. Each simulation starts with a single cell of the size 15 *×* 15 pixels placed on a substrate of width of 450 pixels and allowed to proliferate following Equation (3). The simulation domain is surrounded by wall cells that prevent the cells from sticking to the lattice boundaries. The wall cells are excluded from participating in the pixel copies of the Potts model [39]. If a cell division occurs, the cell is divided along a plane specified by a normal vector *n*_div_ = (*n*_*x*_, *n*_*z*_), where *n*_*x*_ and *n*_*z*_ are the components normal to the plane. The division then results in two cells each with area *≈ A*_0_*/*2. Then according to Equations (1, 2) these two cells grow to reach the target area *A*_0_. Table 1 summarises the parameter values used in computational simulations.

**Table 1:** Model parameters.

| Parameter | Value |
| --- | --- |
| Initial cell size (pixel × pixel) | $15 \times 15$ |
| Initial cell area, $A_\sigma$ (pixel × pixel) | 225 |
| Preferred area, $A_0$ (pixel × pixel) | 225 |
| Temperature, $T$ | 50 |
| Half-saturation constant, $\gamma$ | 2 |
| Hill coefficient, $k$ | 10 |
| Maximal proliferation probability, $P_{\text{max}}$ | 0.1 |

## 3 Results and Discussion

### 3.1 Single cell morphology

Since the multicellular morphogenesis is partly driven by changes in the shape of individual cells [40], our starting point for modelling collective epithelial cell morphology is to explore single-cell morphology in response to its mechanical properties, when cell proliferation is switched off. Typical snapshots of single-cell morphology in the steady-state are shown in Fig. 1. We find that the squamous (i.e., flat)-to-cuboidal shape transition is promoted by increasing cell contractility. A similar shape transition is also found with decreasing cell-substrate adhesion; see Fig. 1(a, b).

**Figure 1:**
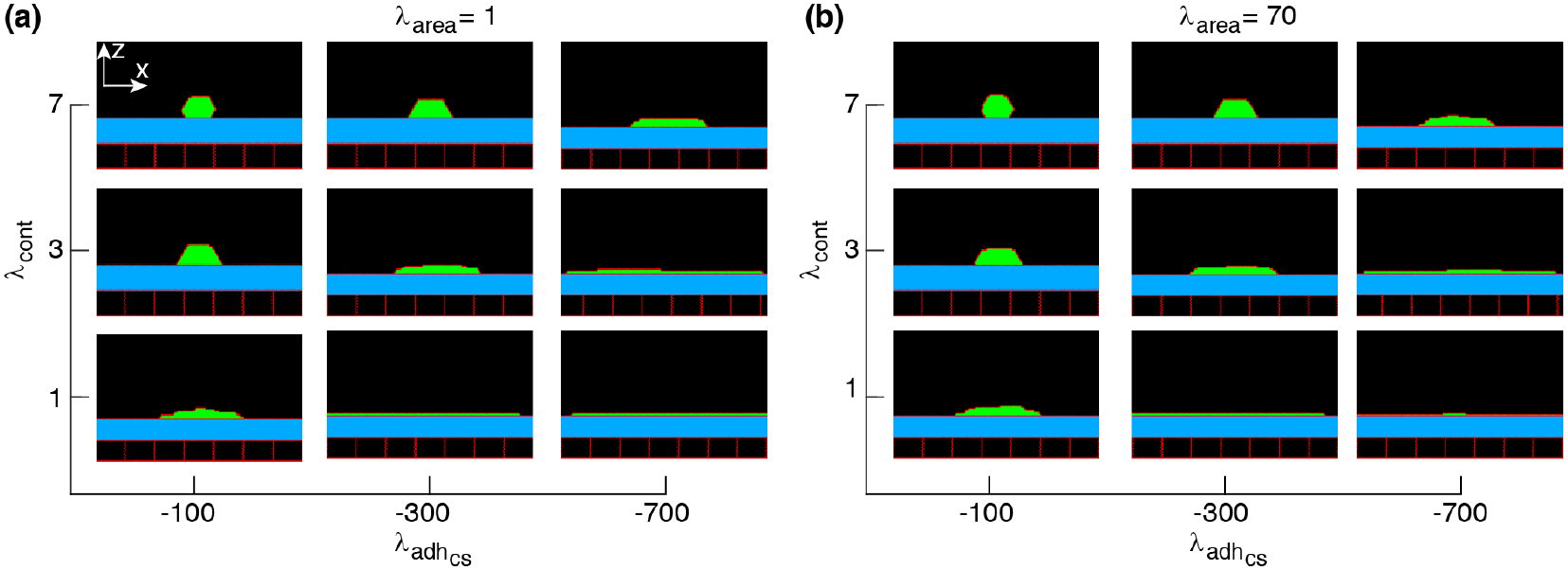
Phase diagram of single-cell shapes. *x*-*z* cross-section of the cell (green) placed on a substrate (blue) surrounded by an empty region (black) and wall cells (black squares). Simulations were run with the cell proliferation disabled. The range of parameter values are adopted from [26]. Snapshots were taken in the steady-states.

To better understand how the single-cell morphology can influence the multi-cellular dynamics, we analyse the cell area and number of cell-substrate adhesion pixel (which affect the probability of cell proliferation) in response to the mechanical control parameters. This enables us to predict the combination of mechanical properties that can lead to different collective cell behaviors.

Figure 2(a) shows that the average cell area increases with cell-substrate adhesion, which is more evident with weak cell contractility. Contrarily, with strengthening cell contractility and decreasing cell-substrate adhesion the average area of a cell falls below its target area *A*_0_. On the other hand, with increasing *λ*_area_, the average cell area remains close to *A*_0_ at all *λ*_cont_ and 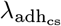; see Fig. 2(b). Further, increasing cell-substrate adhesion and weakening cell contractility expand cell-substrate adhesion sites; see Fig. 2(c, d). Together, these numerical results suggest that monolayers and multilayered structures are more likely to form with increasing cell-substrate adhesion and weaker cell contractility, due to increased proliferation probability of individual cells. Further, non-confluent structures are generated when cells have strong cortex contractility and low adhesion to the substrate.

**Figure 2:**
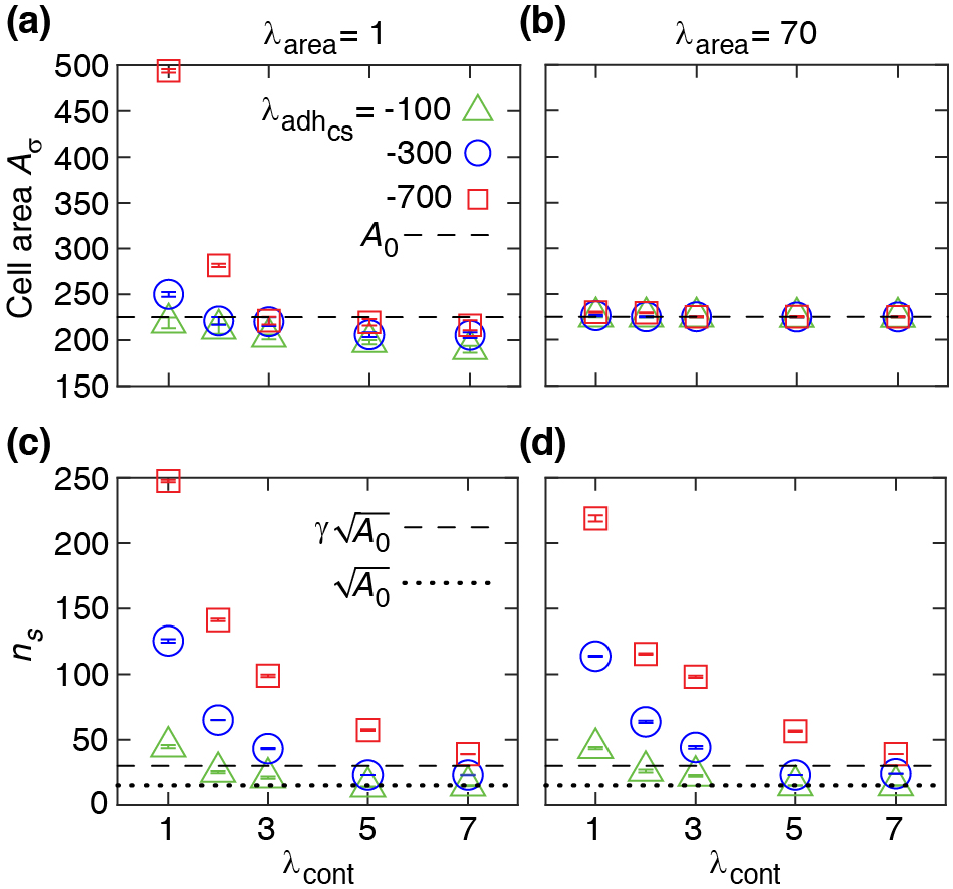
Numerical results for single-cell properties. (a, b) Area of the cell in steady state versus contractility strength *λ*_cont_ at various *λ*_area_ and cell-substrate adhesion 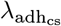. (c, d) Number of cell-substrate adhesion sites *n*_*s*_ in the steady state versus *λ*_cont_ at various *λ*_area_ and 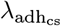. Each symbol is derived from an individual simulation run and corresponds to mean *±* SD.

We check the consistency of these CPM simulation results with the estimated energy minimum determined for a simplified rectangular cell shape. The energy function for a single rectangular cell reads:

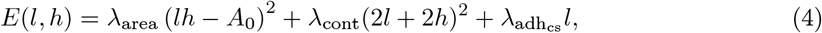

where *l* and *h* are cell length and height (see schematic inset, Fig. 3). The minimum of the energy *E* is determined by solving the equations:

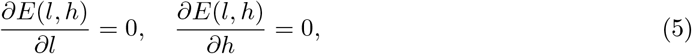

**Figure 3:**
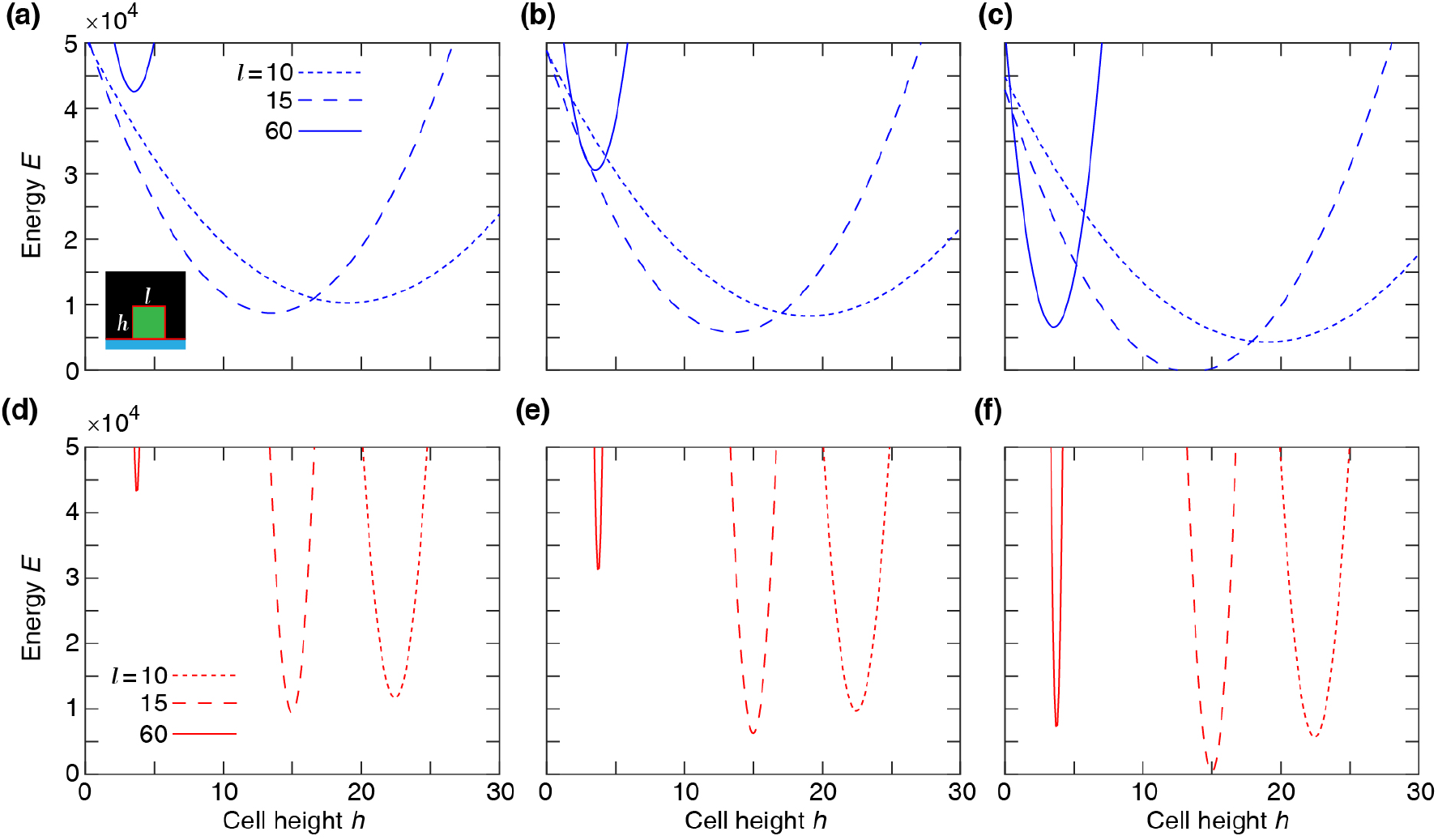
Single-cell Potts energy *E* versus cell height *h* at various cell length *l* calculated using Equation (4). Top row: *λ*_area_ = 1 and *λ*_cont_ = 3 at 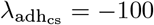 (a), −300 (b), −700 (c). Bottom row: *λ*_area_ = 70 and *λ*_cont_ = 3 at 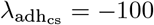 (d), −300 (e), −700 (f).

which results in

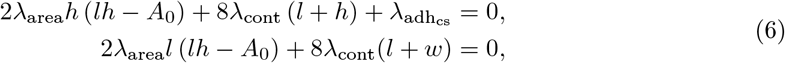

to define cell length *l*^∗^ and height *h*^∗^ at mechanical equilibrium. The typical dependence of the energy function on the cell height and width is illustrated in Fig. 3. Assuming that the mechanical equilibrium at steady state can be approximately estimated from the minimisation of the energy function corresponding to a rectangular cell (4), we calculate the cell aspect ratio and area. The results shown in Fig. 4 are consistent with the phase diagram of single-cell morphology in Fig. 1 and also show qualitative agreement with the CPM simulation results in Fig. 2(a, b).

**Figure 4:**
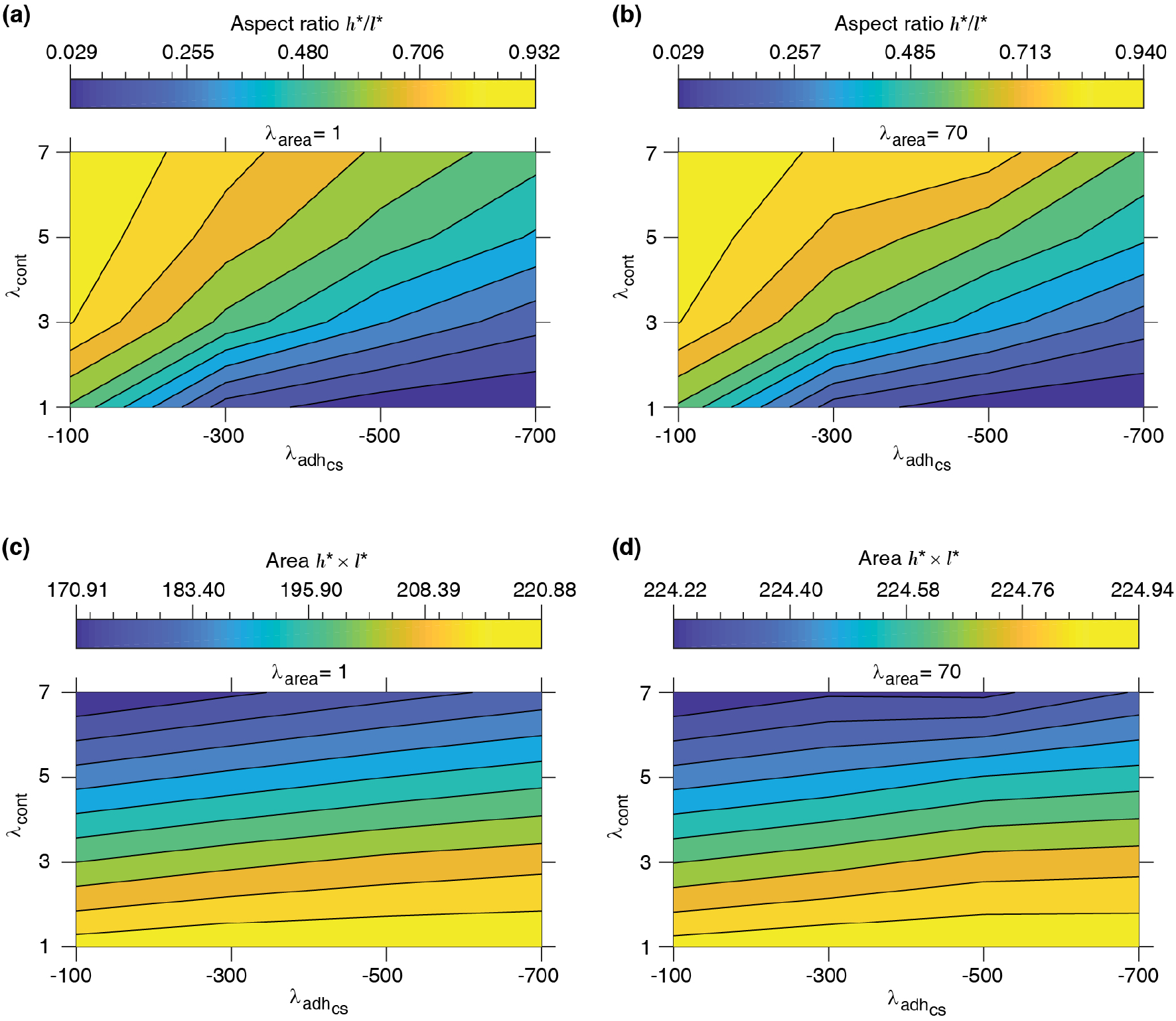
Equilibrium single-cell aspect ratio *w*^∗^*/L*^∗^ and area *w*^∗^ *× L*^∗^ versus cell-substrate adhesion 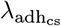 and contractility *λ*_cont_. *l*^∗^ and *h*^∗^: cell length and height at the mechanical equilibrium, respectively, corresponding to rectangular cell shape; see Equation (4).

### 3.2 Collective multicellular morphology

We now use the model to simulate a system of proliferating cells using various combinations of mechanical parameters. The simulations are started with a single cell placed on the substrate in the middle of the domain and cell division is allowed according to the rules described above, i.e. when the cell area is larger than *A*_0_ with a probability dependent on the number cell adhesion sites (pixels) attached to the substrate.

During morphogenesis, oriented cell divisions are essential for the generation of cell diversity and for tissue shaping [11]. Yet, current evidence on the role of cell shape and different sets of intracellular mechanisms in orienting cell proliferation remains inconclusive [11, 12]. Our model allows a convenient way to simulate collective cell morphologies by considering different orientations of cell division axis and varying mechanical properties of cells in various combinations.

Steady state phase diagrams of the collective morphology with horizontal, vertical, and random cell division orientation are presented in Figs. 5-7. First, we note that the cell shapes in the multicellular system in most cases can be quite different from the shape of a single isolated cell obtained for the same set of mechanical parameters. We observe three main types of multi-cellular structures and behaviors developing in the simulations: 1) For certain parameter combinations, the cell division is either completely blocked or is very limited resulting in the formation of a small group of cells without forming a confluent cell layer along the substrate over the whole domain. 2) A cell monolayer can form through repeated cell divisions in such a way that cell proliferation stops in a self-regulated manner once a fully confluent layer is formed. This layer may be composed of flat or tall cells. 3) In multi-layered structures, the cell division continues indefinitely (although it is still restricted to the basal cells along the substrate) and the height of the cell layer increases over time. In a real multilayered epithelium, the height of such layer can be controlled by differentiation and death of the non-proliferating cells that are not adhered to the substrate. Since we are focusing on the emergence of the different cell layer structures and the corresponding cell shapes, we do not include cell death and differentiation in our model.

**Figure 5:**
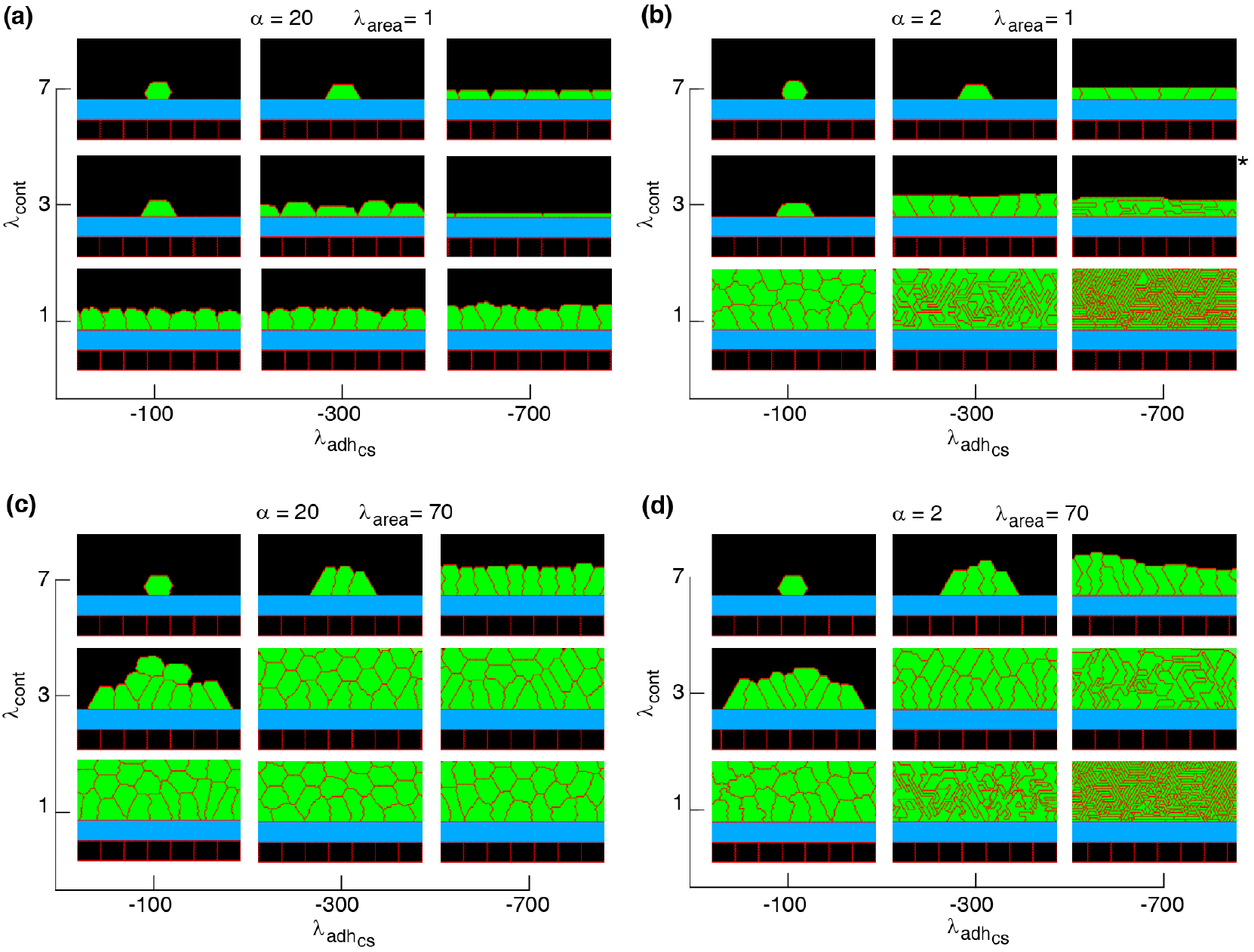
Phase diagram of steady state collective cell morphology when the axis of division is perpendicular to the substrate, i.e., *n*_div_ = (1, 0); see Equation (3). 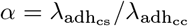. See Movies 1-5.

**Figure 6:**
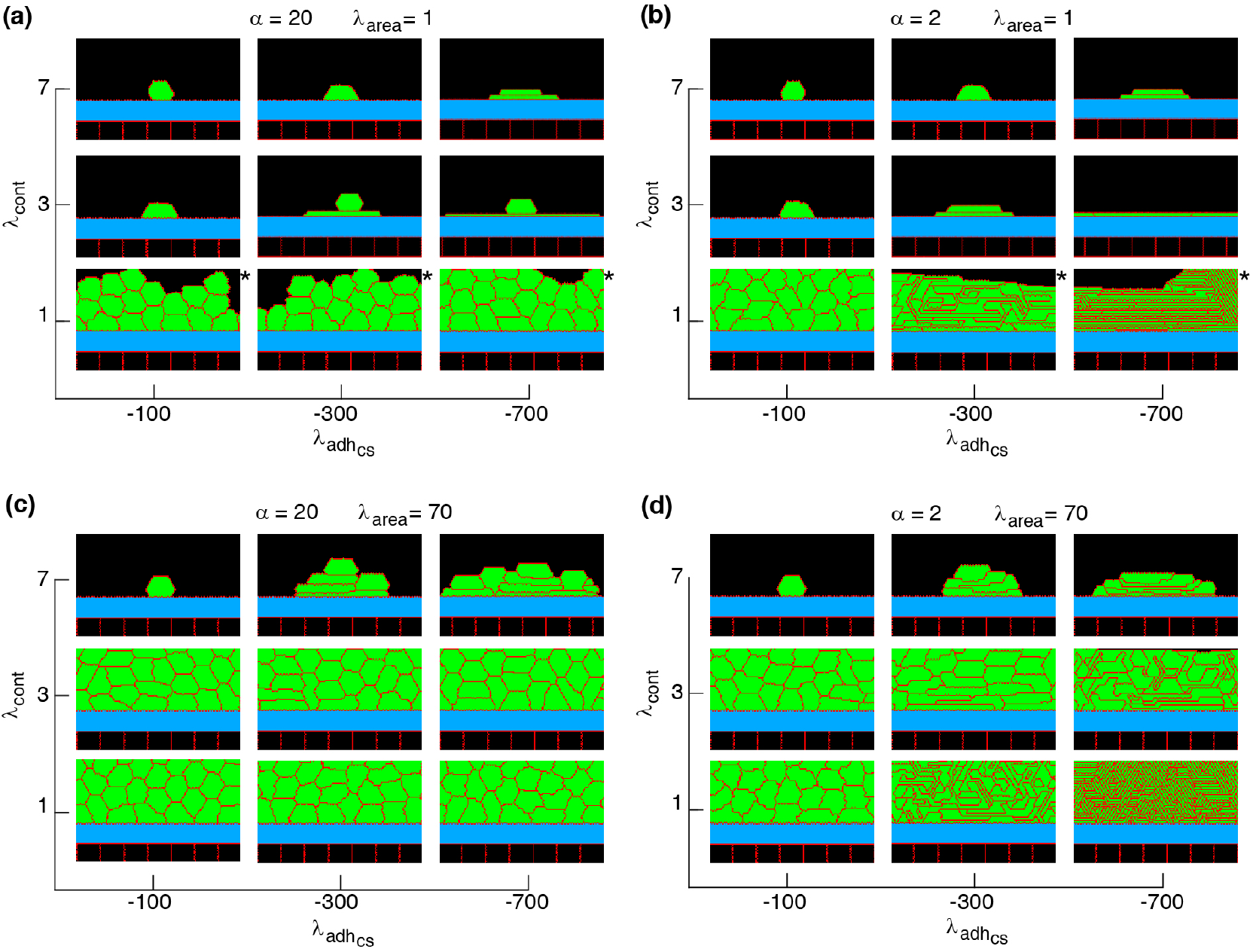
Phase diagram of collective cell morphology with horizontal orientation of cell proliferation, where the axis of division is parallel to the substrate, i.e., *n*_div_ = (0, 1); see Equation (3). 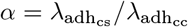. See Movies 6, 7.

**Figure 7:**
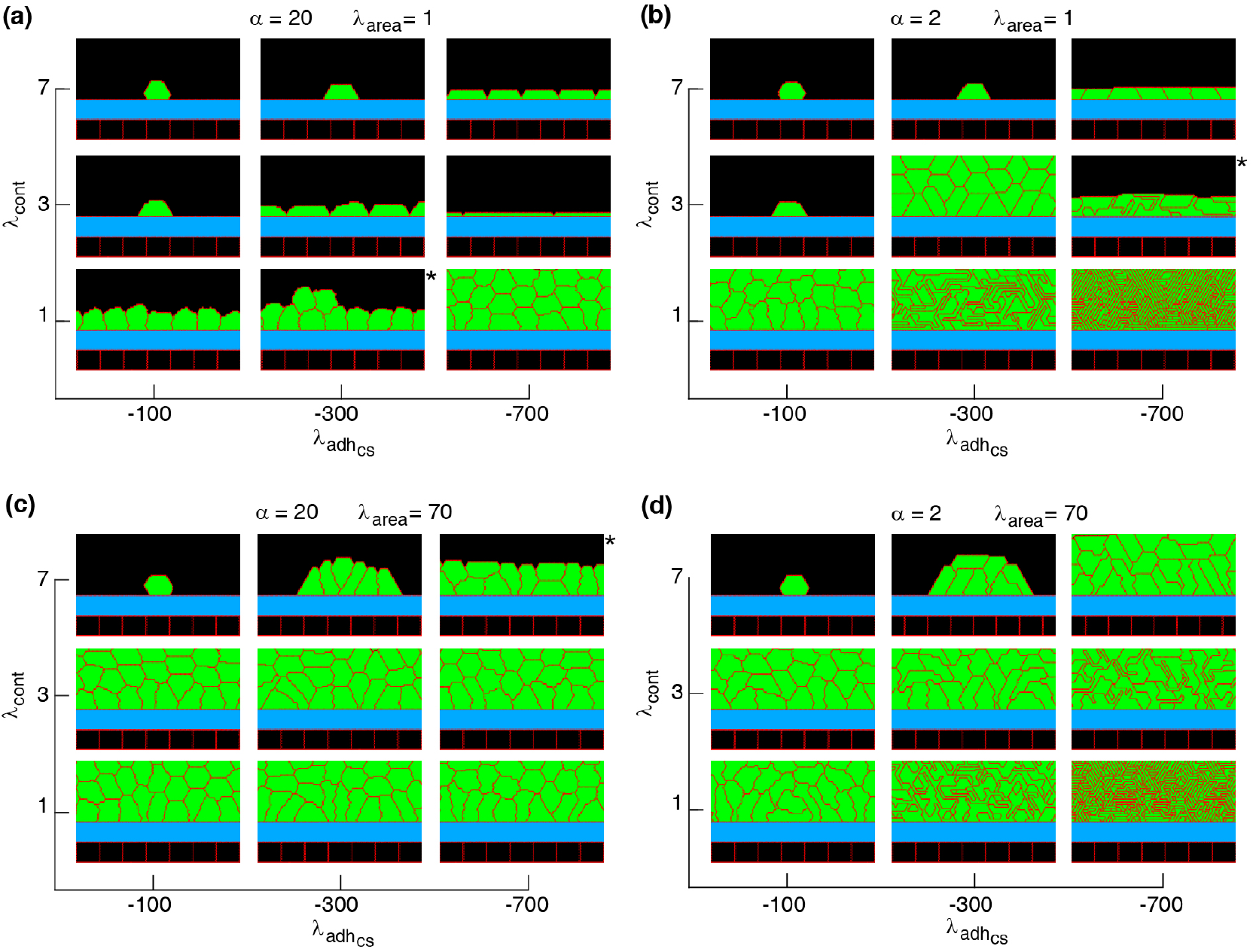
Phase diagram of collective cell morphologies with random orientation of cell proliferation. 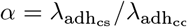. See Movies 8, 9.

Non-confluent structures are formed at high cell contractility and reduced cell-substrate adhesion, independently of the proliferation orientation; see Figs. 5-7. This is due to reductions in both cell area and the number of cell-substrate adhesion sites in individual cells which reduce the probability of cell proliferation; see Fig. 2.

Confluent monolayers and multilayered structures are formed with increasing cell-substrate adhesion and lowering cell contractility. With vertical proliferation orientation, monolayers of squamous (flat), cuboidal, and columnar (tall) cells are found; see Fig. 5. The expansion of monolayers typically happens through the division of border cells (at both edges of the monolayer) on the substrate, while the other cells inside the monolayer do not divide or only relatively rarely. Once the layer becomes confluent cell crowding limits the cell area and the cell-substrate adhesion sites, due to which the probability of proliferation decreases; see Fig. 8(a-i). At certain combinations of mechanical properties (summarised in Fig. 5 and analysed in Fig. 8(a-i)), cells stop proliferating once a confluent monolayer is formedsee Movies 1-3, 5. Squamous cells are mostly found for high cell-substrate adhesion (relative to cell-cell adhesion), increased cortex contractility and reduced *λ*_area_ parameter. Increasing *λ*_area_, in this regime, leads to squamous-to-columnar shape transition.

**Figure 8:**
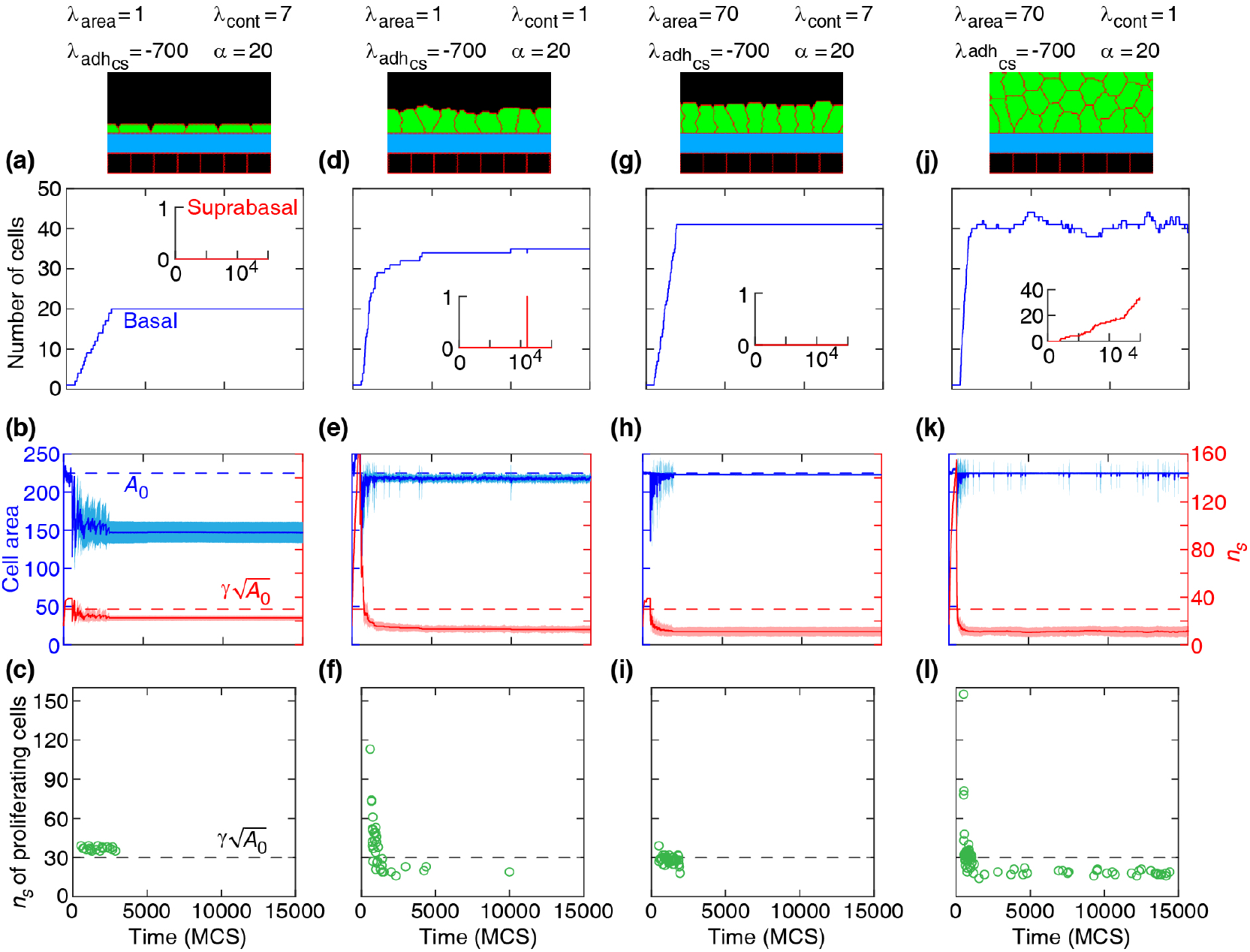
Dynamics of basal cells in four different collective morphologies illustrated with snapshots and associated mechanical parameters. Top row: number of basal cells (with adherence to the substrate) and suprabasal cells (without adherence to the substrate) versus time. Middle row: area (left axis) and number of cell-substrate adhesion sites *n*_*s*_ (right axis) for basal cells versus time. Solid curves: mean. Shaded region: SD. Bottom row: *n*_*s*_ of proliferating cells versus time. Orientation of cell proliferation is vertical, *n*_div_ = (1, 0). 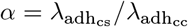. See Movies 1-4.

A major factor that contributes to monolayer-to-multilayer transition is cell crowding. When the cell density cannot increase anymore in the basal layer, while cell deformations are likely, cells are extruded from the monolayer. Accordingly, cells at the basal layer can expand their area increasing the probability of their proliferation and further extrusion events; see Fig. 8(j-l) and Movie 4. Overall, monolayer-to-multilayer transition is more likely to appear with a combination of parameters that increase *λ*_area_, increasing cell-substrate adhesion, reduced cortical contractility, and with random proliferation orientation or being parallel to the substrate; see Figs. 6 and 7.

Our simulation results complement earlier findings on collective epithelial morphology. Simulation results characterise cell area strength *λ*_area_ as a major factor that influences collective morphology: increasing *λ*_area_ generates multilayer structures, whereas with reducing *λ*_area_ monolayer and non-confluent structures appear; see Figs. 5-7. Increasing *λ*_area_ promotes proliferation probability, by increasing cell area to approach the target area *A*_0_; see Equation 1. At a low *λ*_area_, it is more likely that cell area deviates from *A*_0_ resulting in smaller probability of proliferation. This effect is evident in the generation of monolayers, where cell proliferation is limited by cell area and the number of cell-substrate adhesion sites; see Fig. 8(a-i). However, in multilayer structures, the growth of cell area is facilitated (by *λ*_area_, *λ*_cont_, and 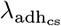) so that the crowding does not block cell proliferation events and continuous cell extrusions out of the basal layer lead to multilayered structure; see Fig. 8(j-l). At stronger cell-substrate adhesion, the extruded cells may return back to the basal layer; see inset in Fig. 8(d).

These simulation results are consistent with experimental observations. It has been observed that the probability of cell proliferation increases with cell area [5] and reduction in cell area (imposed by mechanical constraints on tissue expansion) inhibits cell proliferation [4, 41]. Further, substrate stiffness has been known to be positively correlated with cell proliferation increasing substrate stiffness (dependent on cell-substrate adhesion) and was found to increase the proliferation rate [9, 10, 42]. It was shown that when the cell density cannot increase anymore in a monolayer (due to cell crowding), while the proliferation events still occur, newly generated cells are extruded out of the monolayer where they remained without adhering to the substrate [43]. These suprabasal cells may slide over the basal cells such that they migrate in and become basal cells themselves [44, 45].

Our simulations of cell shapes in monolayers are also consistent with experimental observations. For example, with intermediate cortical contractility *λ*_cont_ = 3 and 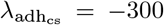, increasing adhesion ratio *α* from 2 to 20 reduces cell height by factor 1.88 ; compare middle snapshots in Fig. 5(a, b). For squamous cells (*λ*_cont_ = 7 and 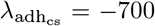), cell height drops by factor 1.22; compare top right corner snapshots in Fig. 5(a, b). For columnar cells, a negligible reduction (by factor 1.10) in cell height is found with increasing *α*; compare top right corner snapshots in Fig. 5(c, d). These simulation results agree with experimental observations that lowering the lateral cell–cell adhesion decreases cell height [46–49]. It is also consistent with the theoretical prediction that the cell-cell lateral adhesion is a crucial parameter to increase cell height [1, 50]. Further, the thin multicellular shapes, formed at high adhesion and low contractility, are consistent with the so called “soft” cell shape regime [17, 26, 31].

## 4 Conclusion

In this article, we introduced a minimal computational model to investigate the emergence of collective morphology of epithelial cells. The model allowed us to simulate diverse collective morphology using various combinations of mechanical properties of cells and the orientation of cell division axis. Our results suggest that non-confluent structures transition into confluent monolayers and multilayers with weakening cell contractility (*λ*_cont_) and strengthening cell-substrate adhesion 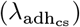, due to increase in probability of cell proliferation. Confluent monolayers of squamous, cuboidal, and columnar cells are formed with proliferation axis perpendicular to the substrate. It is further suggested that monolayer-to-multilayer transition occurs by cell extrusion from the basal layer as a result of the interplay between mechanical parameters (*λ*_area_, *λ*_cont_, and 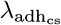) and the orientation of cell proliferation. Taken together, our simulation results suggest that desirable biomechanical features of individual cells can regulate multicellular tissue morphology.

## Acknowledgements

H.K and Z.N. was supported by ARC Discovery Project No. DP160104342.

## Supplementary Material

**Movie 1**. A simulation of a monolayer of squamous cells, where the axis of cell proliferation is perpendicular to the substrate, i.e., *n*_div_ = (1, 0); see Equation (3). Simulation parameters are: *λ*_area_ = 1, *λ*_cont_ = 7, 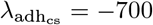, and 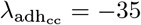.

**Movie 2**. A simulation of a monolayer of cuboidal cells, where the axis of cell proliferation is perpendicular to the substrate, i.e., *n*_div_ = (1, 0); see Equation (3). Simulation parameters are: *λ*_area_ = 1, *λ*_cont_ = 1, 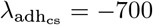, and 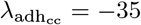.

**Movie 3**. A simulation of a monolayer of columnar cells. Simulation parameters are: *λ*_area_ = 70, *λ*_cont_ = 7, 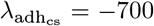, and 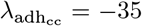. The axis of cell proliferation is perpendicular to the substrate, i.e., *n*_div_ = (1, 0); see Equation (3).

**Movie 4**. A simulation of a multilayer structure, where the axis of cell proliferation is perpendicular to the substrate, i.e., *n*_div_ = (1, 0); see Equation (3). Simulation parameters are: *λ*_area_ = 70, *λ*_cont_ = 1, 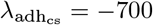, and 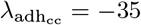.

**Movie 5**. A simulation of a monolayer of cuboidal cells, where the axis of cell proliferation is perpendicular to the substrate, i.e., *n*_div_ = (1, 0); see Equation (3). Simulation parameters are: *λ*_area_ = 1, *λ*_cont_ = 1, 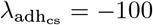, and 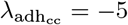.

**Movie 6**. A simulation of a non-confluent structure. Simulation parameters are: *λ*_area_ = 1, *λ*_cont_ = 3, 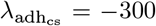, and 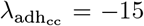. The axis of cell proliferation is parallel to the substrate, i.e., *n*_div_ = (0, 1); see Equation (3).

**Movie 7**. A simulation of a multilayer structure, where the axis of cell proliferation is parallel to the substrate, i.e., *n*_div_ = (0, 1); see Equation (3). Simulation parameters are: *λ*_area_ = 70, *λ*_cont_ = 3, 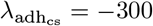, and 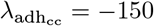.

**Movie 8**. A simulation of a monolayer of squamous cells, with random orientation of cell proliferation. Simulation parameters are: *λ*_area_ = 1, *λ*_cont_ = 3, 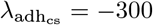, and 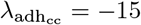.

**Movie 9**. A simulation of a multilayer structure, with random orientation of cell proliferation. Simulation parameters are: *λ*_area_ = 70, *λ*_cont_ = 3, 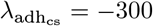, and 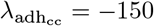.

## References

1 E. Hannezo, J. Prost, and J.-F. Joanny, “Theory of epithelial sheet morphology in three dimensions”, Proceedings of the National Academy of Sciences 111, 27–32 (2014).

2 R. Vincent, E. Bazellières, C. Pérez-González, M. Uroz, X. Serra-Picamal, and X. Trepat, “Active Tensile Modulus of an Epithelial Monolayer”, Physical Review Letters 115, 248103 (2015).

3 S. F. Pedersen, E. K. Hoffmann, and I. Novak, “Cell volume regulation in epithelial physiology and cancer”, Frontiers in Physiology 4, 1–12 (2013).

4 C. S. Chen, “Geometric control of cell life and death”, Science 276, 1425–1428 (1997).

5 S. J. Streichan, C. R. Hoerner, T. Schneidt, D. Holzer, and L. Hufnagel, “Spatial constraints control cell proliferation in tissues”, Proceedings of the National Academy of Sciences 111, 5586– 5591 (2014).

6 M. A. Schwartz and R. K. Assoian, “Integrins and cell proliferation: regulation of cyclindependent kinases via cytoplasmic signaling pathways”, Journal of Cell Science 114, 2553–2560 (2001).

7 C. Brakebusch, D. Bouvard, F. Stanchi, T. Sakai, and R. Fässler, “Integrins in invasive growth”, Journal of Clinical Investigation 109, 999–1006 (2002).

8 S. A. Hacking and A. Khademhosseini, “Cells and surfaces in vitro”, in Biomaterials science an introduction to materials in medicine, edited by B. D. Ratner, A. S. Hoffman, F. J. Schoen, and J. E. Lemons (Elsevier, 2013), pp. 408–427.

9 P. P. Provenzano and P. J. Keely, “Mechanical signaling through the cytoskeleton regulates cell proliferation by coordinated focal adhesion and Rho GTPase signaling”, Journal of Cell Science 124, 1195–1205 (2011).

10 A. Mohan, K. T. Schlue, A. F. Kniffin, C. R. Mayer, A. A. Duke, V. Narayanan, P. T. Arsenovic, K. Bathula, B. E. Danielsson, S. P. Dumbali, V. Maruthamuthu, and D. E. Conway, “Spatial Proliferation of Epithelial Cells Is Regulated by E-Cadherin Force”, Biophysical Journal 115, 853–864 (2018).

11 T. M. Finegan and D. T. Bergstralh, “Division orientation: disentangling shape and mechanical forces”, Cell Cycle 18, 1187–1198 (2019).

12 C. Collinet and T. Lecuit, “Stability and dynamics of cell-cell junctions”, Progress in Molecular Biology and Translational Science 116, 25–47 (2013).

13 A. Hočevar Brezavšček, M. Rauzi, M. Leptin, and P. Ziherl, “A Model of Epithelial Invagination Driven by Collective Mechanics of Identical Cells”, Biophysical Journal 103, 1069–1077 (2012).

14 M. Krajnc, N. Štorgel, A.H. Brezavšček, and P. Ziherl, “A tension-based model of flat and corrugated simple epithelia”, Soft Matter 9, 8368 (2013).

15 J. Kafer, T. Hayashi, A. F. M. Maree, R. W. Carthew, and F. Graner, “Cell adhesion and cortex contractility determine cell patterning in the Drosophilaretina”, Proceedings of the National Academy of Sciences 104, 18549–18554 (2007).

16 S. Hilgenfeldt, S. Erisken, and R. W. Carthew, “Physical modeling of cell geometric order in an epithelial tissue”, Proceedings of the National Academy of Sciences 105, 907–911 (2008).

17 R. Farhadifar, J.-C. Röper, B. Aigouy, S. Eaton, and F. Jülicher, “The Influence of Cell Mechanics, Cell-Cell Interactions, and Proliferation on Epithelial Packing”, Current Biology 17, 2095– 2104 (2007).

18 M. C. Gibson, A. B. Patel, R. Nagpal, and N. Perrimon, “The emergence of geometric order in proliferating metazoan epithelia”, Nature 442, 1038–1041 (2006).

19 M. Osterfield, X. Du, T. Schüpbach, E. Wieschaus, and S. Y. Shvartsman, “Three-Dimensional Epithelial Morphogenesis in the Developing Drosophila Egg”, Developmental Cell 24, 400–410 (2013).

20 V. Kostiou, M. W. J. Hall, P. H. Jones, and B. A. Hall, “Different responses to cell crowding determine the clonal fitness of p53 and notch inhibiting mutations in squamous epithelia”, bioRxiv, 1–25 (2020).

21 D.P. Doupé, A. M. Klein, B. D. Simons, and P. H. Jones, “The Ordered Architecture of Murine Ear Epidermis Is Maintained by Progenitor Cells with Random Fate”, Developmental Cell 18, 317–323 (2010).

22 D. P. Doupe, M. P. Alcolea, A. Roshan, G. Zhang, A. M. Klein, B. D. Simons, and P. H. Jones, “A Single Progenitor Population Switches Behavior to Maintain and Repair Esophageal Epithelium”, Science 337, 1091–1093 (2012).

23 F. Graner and J. A. Glazier, “Simulation of biological cell sorting using a two-dimensional extended potts model”, Physical Review Letters 69, 2013–2016 (1992).

24 J. A. Glazier and F. Graner, “Simulation of the differential adhesion driven rearrangement of biological cells”, Physical Review E 47, 2128–2154 (1993).

25 F. Kempf, A. Goychuk, and E. Frey, “Tissue flow through pores: a computational study”, bioRxiv, 1–29 (2021).

26 H. Khataee, A. Czirok, and Z. Neufeld, “Multiscale modelling of motility wave propagation in cell migration”, Scientific Reports 10, 8128 (2020).

27 J. M. Osborne, A. G. Fletcher, J. M. Pitt-Francis, P. K. Maini, and D. J. Gavaghan, “Comparing individual-based approaches to modelling the self-organization of multicellular tissues”, PLOS Computational Biology 13, e1005387 (2017).

28 R. Giniūnaitė, R. E. Baker, P. M. Kulesa, and P. K. Maini, “Modelling collective cell migration: neural crest as a model paradigm”, Journal of Mathematical Biology (2019).

29 F. Zhou, S. A. Schaffer, C. Schreiber, F. J. Segerer, A. Goychuk, E. Frey, and J.O. Rädler, “Quasi-periodic migration of single cells on short microlanes”, PLOS ONE 15, e0230679 (2020).

30 F. J. Segerer, F. Thüroff, A. Piera Alberola, E. Frey, and J.O. Rädler, “Emergence and Persistence of Collective Cell Migration on Small Circular Micropatterns”, en, Physical Review Letters 114, 228102 (2015).

31 A. R. Noppe, A. P. Roberts, A. S. Yap, G. A. Gomez, and Z. Neufeld, “Modelling wound closure in an epithelial cell sheet using the cellular potts model”, Integrative Biology 7, 1253–1264 (2015).

32 P. J. Albert and U. S. Schwarz, “Dynamics of cell ensembles on adhesive micropatterns: bridging the gap between single cell spreading and collective cell migration”, PLOS Computational Biology 12, e1004863 (2016).

33 F. Thüroff, A. Goychuk, M. Reiter, and E. Frey, “Bridging the gap between single-cell migration and collective dynamics”, eLife 8, e46842 (2019).

34 M. Reffay, M. C. Parrini, O. Cochet-Escartin, B. Ladoux, A. Buguin, S. Coscoy, F. Amblard, J. Camonis, and P. Silberzan, “Interplay of rhoa and mechanical forces in collective cell migration driven by leader cells”, Nature Cell Biology 16, 217–223 (2014).

35 G. Charras and A. S. Yap, “Tensile forces and mechanotransduction at cell–cell junctions.”, Current Biology 28, R445–R457 (2018).

36 M. H. Swat, G. L. Thomas, J. M. Belmonte, A. Shirinifard, D. Hmeljak, and J. A. Glazier, “Multiscale modeling of tissues using compucell3d”, Methods in cell Biology 110, 325–366 (2012).

37 M. Santillán, “On the Use of the Hill Functions in Mathematical Models of Gene Regulatory Networks”, Mathematical Modelling of Natural Phenomena 3, 85–97 (2008).

38 N. Paddillaya, A. Mishra, P. Kondaiah, P. Pullarkat, G. I. Menon, and N. Gundiah, “Biophysics of Cell-Substrate Interactions Under Shear”, Frontiers in Cell and Developmental Biology 7, 251 (2019).

39 M. H. Swat, S. D. Hester, A. I. Balter, R. W. Heiland, B. L. Zaitlen, and J. A. Glazier, “Multi-cell simulations of development and disease using the compucell3d simulation environment”, Methods in molecular biology 500, 361–428.

40 T. J. Widmann and C. Dahmann, “Dpp signaling promotes the cuboidal-to-columnar shape transition of Drosophila wing disc epithelia by regulating Rho1”, Journal of Cell Science 122, 1362–1373 (2009).

41 A. Puliafito, L. Hufnagel, P. Neveu, S. Streichan, A. Sigal, D. K. Fygenson, and B. I. Shraiman, “Collective and single cell behavior in epithelial contact inhibition”, Proceedings of the National Academy of Sciences 109, 739–744 (2012).

42 A. Garcia, A. Vega, and D. Boettiger, “Modulation of Cell Proliferation and Differentiation through Substrate-dependent Changes in Fibronectin Conformation”, Molecular Biology of the Cell 10, 785–798 (1999).

43 M. Deforet, V. Hakim, H. Yevick, G. Duclos, and P. Silberzan, “Emergence of collective modes and tri-dimensional structures from epithelial confinement”, Nature Communications 5, 3747 (2014).

44 E. Rognoni and F. M. Watt, “Skin Cell Heterogeneity in Development, Wound Healing, and Cancer”, Trends in Cell Biology 28, 709–722 (2018).

45 D. Haensel, S. Jin, P. Sun, R. Cinco, M. Dragan, Q. Nguyen, Z. Cang, Y. Gong, R. Vu, A. L. MacLean, K. Kessenbrock, E. Gratton, Q. Nie, and X. Dai, “Defining Epidermal Basal Cell States during Skin Homeostasis and Wound Healing Using Single-Cell Transcriptomics”, Cell Reports 30, 3932–3947.e6 (2020).

46 K. L. Weber, R. S. Fischer, and V. M. Fowler, “Tmod3 regulates polarized epithelial cell morphology”, Journal of Cell Science 120, 3625–3632 (2007).

47 M. Melani, K. J. Simpson, J. S. Brugge, and D. Montell, “Regulation of Cell Adhesion and Collective Cell Migration by Hindsight and Its Human Homolog RREB1”, Current Biology 18, 532–537 (2008).

48 J. M. Gomez, Y. Wang, and V. Riechmann, “Tao controls epithelial morphogenesis by promoting Fasciclin 2 endocytosis”, Journal of Cell Biology 199, 1131–1143 (2012).

49 D. J. Montell, “Morphogenetic Cell Movements: Diversity from Modular Mechanical Properties”, Science 322, 1502–1505 (2008).

50 K. Dasbiswas, E. Hannezo, and N. S. Gov, “Theory of Epithelial Cell Shape Transitions Induced by Mechanoactive Chemical Gradients”, Biophysical Journal 114, 968–977 (2018).

